# Population Differentiation at the *HLA* Genes

**DOI:** 10.1101/214668

**Authors:** Débora Y. C. Brandt, Jônatas César, Jérôme Goudet, Diogo Meyer

## Abstract

Balancing selection is defined as a class of selective regimes that maintain polymorphism above what is expected under neutrality. Theory predicts that balancing selection reduces population differentiation, as measured by F_ST_. However, balancing selection regimes in which different sets of alleles are maintained in different populations could increase population differentiation. To tackle this issue, we investigated population differentiation at the HLA genes, which constitute the most striking example of balancing selection in humans. We found that population differentiation of single nucleotide polymorphisms (SNPs) at the HLA genes is on average lower than that of SNPs in other genomic regions. However, this result depends on accounting for the differences in allele frequency between selected and putatively neutral sites. Our finding of reduced differentiation at SNPs within HLA genes suggests a predominant role of shared selective pressures among populations at a global scale. However, in pairs of closely related populations, where genome-wide differentiation is low, differentiation at HLA is higher than in other genomic regions. This pattern was reproduced in simulations of overdominant selection. We conclude that population differentiation at the HLA genes is generally lower than genome-wide, but it may be higher for recently diverged population pairs, and that this pattern can be explained by a simple overdominance regime.

---

Natural selection is one of the forces shaping the genetic variation within and the differentiation between populations. In the case of a locus where a variant is favored in one population but not in another *(i.e.,* in which selection drives local adaptation), we expect differentiation to exceed that under purely demographic processes (Lewontin and Krakauer 1973). This is the case for well known examples, such as the regulatory variant that generates lactase persistence in adulthood, which is more frequent in Europeans (Bersaglieri *et al.* 2004) and variants of the *EPAS* gene that provide adaptation to high altitude in Tibetans (Xu *et al.* 2011). Purifying selection, on the other hand, is more common and removes most variants which would contribute to differences among populations. Therefore, it is expected to reduce genetic differentiation at the focal locus with respect to a strictly neutral scenario (*e.g.* Barreiro *et al.* (2008)), while differentiation in surrounding genomic regions may increase due to the lower effective population size (Charlesworth *et al.* 1997).

A third regime, balancing selection, is related to diversity and differentiation in more complex ways. By definition balancing selection encompasses all selective regimes that result in increased genetic diversity with respect to neutral expectations. The increased variability can result from a variety of processes, often with different underlying biological properties: frequency dependent selection, heterozygote advantage, selection varying over temporal and geographic scales (Andrés 2011). As a consequence, the expectations regarding population differentiation under balancing selection represent a challenging theoretical and empirical question.

Across human populations, the loci with the strongest evidence for balancing selection are the classical HLA class I and II loci (especially the *HLA-A,-B,-C,-DRB1,-DQB1* and *-DQA1* loci), which are the human Major Histocompatibility Complex (MHC) genes. These genes encode proteins that mediate a critical step of the adaptive immune response, which is the binding of peptides for presentation on the surface of the cellular membrane. The HLA-peptide complex is surveyed by T-cell receptors, which may trigger an immune response when a non-self peptide is identified (Klein and Sato 2000). Balancing selection at HLA loci has been strongly supported by a wide variety of methods, with evidence including an excess of alleles at intermediate frequency with respect to neutral expectations (Hedrick and Thomson 1983), higher non-synonymous to synonymous substitution rate (Hughes and Nei 1988) and trans-specific polymorphism (Lawlor *et al.* 1988) (Meyer and Thomson 2001, for a review).

Although balancing selection at HLA genes is well documented, the evidence from most studies is compatible with different mechanisms that are difficult to disentangle: heterozygote advantage (Doherty and Zinkernagel 1975; Takahata and Nei 1990; De Boer *et al.* 2004), frequency dependent selection (Slade and McCallum 1992; Borghans *et al.* 2004) and selection that varies over time and space (Eizaguirre *et al.* 2012) have all been proposed to act on the HLA genes.

Interestingly, these selective regimes are theoretically compatible with both increased or reduced population differentiation. For example, the coevolution between HLA and pathogens could create a mechanism of frequency dependent selection, or rare allele advantage. Under this scenario, rare HLA alleles would be advantageous, since few pathogens would have evolved resistance to them (Meyer and Thomson 2001). Rare allele advantage is expected to increase the effective migration rate: migrants will often be rare in the population they arrive to, and thus will be advantageous and increase in frequency in the new population (Schierup *et al.* 2000; Muirhead 2001). Therefore, this regime of balancing selection is expected to reduce population differentiation.

However, in the case of HLA genes, balancing selection may be population-specific, with the different sets of pathogens in each population selecting locally advantageous HLA variants. Under this scenario we expect an increase in population differentiation. Evidence in support of population-specific pathogen selection for humans comes from the finding that HLA and pathogen diversities across populations are correlated (Prugnolle *et al.* 2005), and from theoretical studies showing that population-specific pathogen selection models of balancing selection provide a better explanation for observed HLA variation than heterozygote advantage (Hedrick 2002; Borghans *et al.* 2004).

Pathogen-driven selection implies that specific HLA alleles are more effective in presenting antigens of certain pathogens, to which the population is exposed. Support for this assumption comes from associations between disease susceptibility, resistance or progression with genetic variation at HLA. For example, variants in *HLA-B* are associated to the progression to clinical disease after HIV infection (The International HIV Controllers Study 2010), variants in *HLA-C, -DP* and -DQ are associated to clearance of hepatitis B virus (Hu *et al.* 2013), alleles in *HLA-DRB1* are associated to susceptibility or resistance to leprosy (Vanderborght *et al.* 2007) and alleles at *HLA-B, -DQ* and *-DR* are associated to resistance to severe malaria (Hill *et al.* 1991).

Previous studies which measured population differentiation at the HLA genes found evidence for both increased or reduced population differentiation. For example, Meyer *et al.* (2006) found no significant difference between differentiation at HLA genes and a set of neutral markers used as a control (microsatellites), while Sanchez-Mazas (2007) found lower differentiation at HLA loci than in their genome-wide control (microsatellites and RFLPs). A limitation of these studies is that differences between the neutral genetic markers and the sequence data used for HLA genes introduce confounding variables, making it difficult to determine the roles of selection or characteristics inherent to the marker *(e.g.* mutation rate and diversity). Another study compared differentiation on markers of the same type (microsatellites) located at HLA genes or near them and those located in other genomic regions, which serve as controls (Nunes 2011). This study found increased differentiation in regions near HLA genes. Nonetheless, some issues remain unresolved: Nunes (2011) was mainly interested in native American populations, and used a limited number of markers. Furthermore, the complexity of the mutational mechanism of microsatellites complicates the interpretation of results.

For non-model organisms a similarly wide array of results have been found, with the MHC region (which contains genes homologous to HLA) showing either equal (Miller *et al.* 2010), higher (Loiseau *et al.* 2009; Oliver *et al.* 2009; Cammen *et al.* 2011) or lower (McCairns *et al.* 2011) differentiation than genome-wide averages. These contrasting results could be due to differences in selective regimes among species, or even to variation in selection among genes within a species (Čížková *et al.* 2011).

In summary, it remains unclear whether balancing selection on HLA genes drives increased differentiation due to selection favoring adaptation to locally occurring pathogens, or whether it results in decreased genetic differentiation due to the maintenance of shared polymorphisms among populations.

Here, we revisit the question of population differentiation at the HLA genes through analysis of data from worldwide human populations. We analyze variation at SNPs, which have the advantage of allowing the use of genomic data as an empirical control for HLA SNPs, assuming similar mutation rates for SNPs in the MHC region and the reminder of the genome. Differently from scans that seek genome-wide significance for specific SNPs, we *a priori* define a set of putatively selected SNPs to be surveyed (those within or close to HLA genes). We relate differences in F_ST_ between HLA and non-HLA SNPs to the degree of polymorphism in each of these groups, drawing on recent findings concerning the constraints imposed by allele frequencies on measures of differentiation. Finally, we perform simulations and find a plausible selective regime that reproduces our results.

## Materials and Methods

### SNP Data

SNP genotypes were acquired from the integrated Variant Call Format (VCF) files from phase 3 of the 1000 Genomes Project (1000G) (The 1000 Genomes Project Consortium 2015), which are available at ftp://ftp-trace.ncbi.nih.gov/1000genomes/ftp/release/20130502/.

This dataset includes variants discovered via high coverage exome targeted resequencing and low coverage whole genome resequencing. Because of the higher coverage in exonic regions, all comparisons between HLA SNPs and non-HLA SNPs (which we treated as a control set) were made within the same functional category *(e.g.* intronic or exonic). To this end each SNP was annotated using ANNOVAR (Wang *et al.* 2010). Our findings for differences in F_ST_ between HLA and non-HLA SNPs were qualitatively the same when using either exonic or intronic regions, so throughout the paper we focus on the results for exonic regions.

The 1000G Phase 3 data contains the genomes of 2504 individuals from 26 populations. After applying filters described in the Extended Materials and Methods section, a total of 2000 individuals in 20 populations were kept (see Table S1).

### Estimation of *F*_*ST*_

Population differentiation was calculated as the proportion of variance in allele frequencies among populations (*a*), relative to the total genetic variance (*a + b + c*, with *b* and *c* referring to the variance components between individuals within populations and between gametes within individuals, respectively):

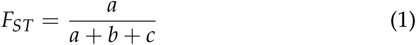

To obtain F_ST_ values we used the Weir and Cockerham (1984) estimator implemented in VCFtools v0.1.14 (Danecek *et al.* 2011). The Weir and Cockerham (1984) estimator was chosen because it is unbiased when sample sizes are large and similar, as in the case of our dataset. F_ST_ was calculated per SNP i) over all populations and ii) for pairs of populations.

When summarizing FST over multiple SNPs, we compared two approaches: i) computing a simple average of F_ST_ at individual SNPs and ii) using the "ratio of averages" approach, suggested by Reynolds *et al.* (1983), in which we first estimate the numerator (*a*) and denominator (*a + b + c*) of F_ST_ for each SNP, and then compute the averages of *a* and *a + b + c* for the desired set of SNPs, and finally compute the ratio of both averages. This second approach provides the least biased estimate of F_ST_, whereas performing a simple average of the F_STS_ of each SNP can lead to an underestimation of differentiation, especially in datasets rich in rare variants (Reynolds *et al.* 1983; Bhatia *et al.* 2013). Unless otherwise stated, we used the "ratio of averages" approach to compute FST. Variance components *(a* and *a* + b + c) were obtained using a minor modification of the VCFtools source code.

### Definition of HLA, peri-HLA and control regions

We define "HLA SNPs" as those contained within the coding sequence of the classical HLA genes *HLA-A,-B,-C,-DRA,-DRB1,-DQA,-DQB1.* Previous studies of *HLA-DPA1* and *-DPB1* found weak or no evidence of balancing selection (Solberg *et al.* 2008; Begovich *et al.* 2001), and even instances of directional selection (Hollenbach *et al.* 2001), making them inappropriate for our question concerning the role of balancing selection on differentiation at HLA loci. Accordingly, our analyses also showed that population differentiation at *HLA-DPA1* and *HLA-DPB1* genes is different from that of other HLA genes (see Results and Figure S1). Therefore, those loci were excluded from our analysis unless otherwise mentioned.

Peri-HLA genes were defined as those that flank the HLA genes and have higher diversity relative to the average of chromosome 6 (Mendes 2013), indicating that their increased polymorphism is driven by hitchhiking to the strongly selected HLA loci (Table S2). These genes are located 119kb to 256kb from the closest HLA locus. All SNPs outside both the HLA and peri-HLA genes comprised the control group.

### Controlling for unreliable allele frequency estimates

The use of a single reference genome to map Next Generation Sequencing (NGS) reads creates mapping bias at some HLA SNPs in the 1000 Genomes Project phase 1 dataset (Brandt *et al.* 2015). We therefore excluded the SNPs within the HLA genes which have unreliable frequency estimates in the 1000 Genomes phase 1 dataset, due to mapping bias (Brandt *et al.* 2015). After applying this filter, 38 out of 525 exonic biallelic SNPs in the HLA genes were excluded (Table S3). In total, 487 SNPs were kept in the HLA group, 1193 in the peri HLA group, and 831,174 in the control group.

In the present study we analyze this filtered version of the 1000 Genomes phase 3 data instead of phase 1 or Sanger sequencing data analyzed in Brandt *et al.* (2015) because it has a larger sample size and includes more populations. Also, SNPs identified as unreliable in the 1000 Genomes phase 1 data were consistent among different populations, which supports the application of this filter to phase 3 data used here.

### Statistical test of F_ST_ differences

When comparing F_ST_ values between HLA and control SNPs, we control for SNPs being located within a small set of genes, resulting in higher linkage disequilibrium (LD) and statistical non-independence than for SNPs in the control group, which comprise a genomewide set. To account for this effect, we designed a strategy to sample the control SNPs so as to approximate the LD structure among the HLA SNPs.

This was done by sampling a random exonic SNP from outside the HLA and peri-HLA genes and selecting all other exonic SNPs from the same gene (resulting in a high LD set of SNPs). We repeated this operation until we obtained a total number of SNPs that matched that of the HLA SNPs, for each MAF bin (see Figure S2). We sampled 1000 such sets of LD-matched control SNPs and compared the F_ST_ distribution of each of those to that of the HLA SNPs, applying a Mann-Whitney test. We recorded the number of comparisons where the difference between F_ST_ distributions was significant (p < 0.05).

### Haplotype level analyses

In addition to SNP based analyses, we also investigated population differentiation when alleles are defined by the coding sequence of each HLA gene (classically referred to as an "HLA allele"). We treat these analyses as "haplotype level", where haplotypes are defined by a combination of SNPs along an HLA gene (i.e., each allele in these analyses is an intragenic haplotype).

Phasing of SNPs in extremely variable regions like the HLA is very challenging, due to the high SNP density and polymorphism. Therefore, rather than estimate intragenic haplotypes directly from the SNP data, we used a publicly available dataset which provides HLA allele calls for samples in the 1000G data based on Sanger sequencing Gourraud *et al.* (2014), available at the dbMHC website (http://www.ncbi.nlm.nih.gov/gv/mhc/xslcgi.fcgi?cmd=cellsearch).

We restricted these haplotype level analyses to *HLA-A*, *-B*, *-C*, *HLA-DRB1* and *-DQB1,* which are reported in Gourraud *et al.* (2014). HLA allele calls were coded so as to only distinguish alleles with nonsynonymous differences *(i.e.,* only the first two fields of the allele names were used, as described in the HLA nomenclature system) (Marsh *et al.* (2010)).

F_ST_ values of multiallelic HLA haplotypes can't be directly compared to those of biallelic SNPs because multiallelic loci tend to have lower allele frequencies, which constrains the maximum value of F_ST_ (Jakobsson *et al.* 2013). To allow F_ST_ values at the HLA haplotypes to be compared to this null distribution, we recoded each HLA gene as a series of biallelic loci. This recoding was done by treating each allele at each gene as "allele 1", and all other alleles as "allele 2". The Weir and Cockerham (1984) F_ST_ estimator was then computed as described for SNPs, using the *wc* function of the hierfstat R package (Goudet 2005).

## Results

### Higher F_ST_ in HLA genes

Initially we compared the distribution of F_ST_ among SNPs from the HLA, peri-HLA and control groups (Figure 1).

**Figure 1.**
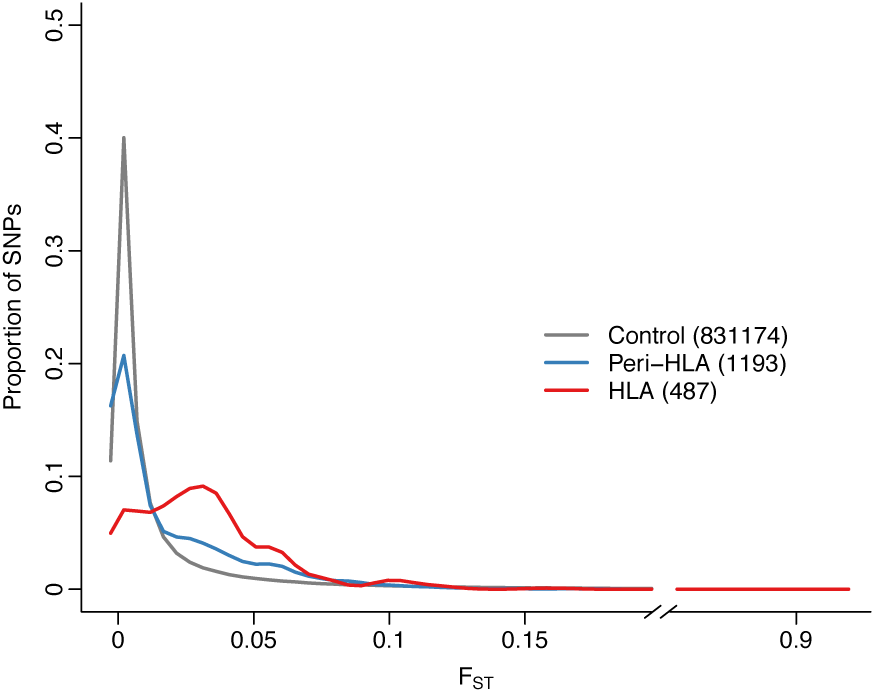
Distribution of F_ST_ values for exonic SNPs from the following groups: outside HLA and peri-HLA regions (control group), within HLA genes and within peri-HLA genes. Number of SNPs in each group is shown in parentheses.

The distribution of F_ST_ of SNPs within HLA genes is shifted towards higher values (median=0.027) compared to control SNPs (median=9.2 * 10^-5^), and the distributions are significantly different (p-value < 10^-16^, two-tailed Mann-Whitney test).

Theory predicts that balancing selection affects only a narrow genomic region, the size of which is defined by the intensity of selection and the recombination rate (Charlesworth *et al.* 1997). To evaluate if the increased differentiation at coding SNPs within HLA genes was also observed in loci that flank the HLA, we applied the same test to the peri-HLA genes. As is the case for HLA SNPs, peri-HLA SNPs have an F_ST_ distribution which is significantly shifted to higher values (median = 0.006, p-value < 10^-16^, two-tailed Mann-Whitney test; Figure 1).

### Lower F_ST_ in HLA when accounting for MAF

The effect of balancing selection is to shift the site frequency spectrum (SFS) of selected loci toward an excess of intermediate frequency variants. This is precisely what we see in the data, with the SFS for HLA SNPs showing a shift to intermediate frequencies compared to control SNPs. The peri-HLA SNPs occupy an intermediate position in the SFS (Figure 2).

**Figure 2.**
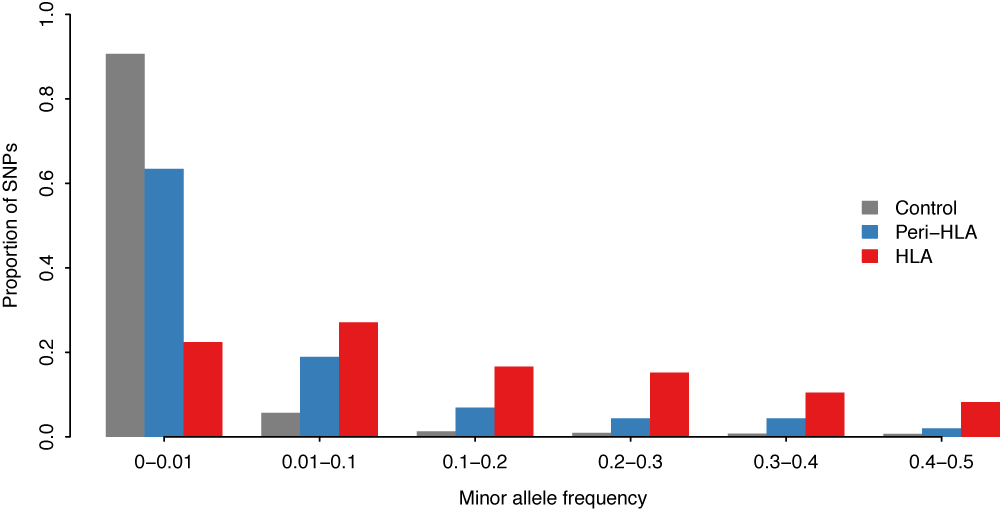
Distribution of minor allele frequency (MAF) for exonic control, peri-HLA genes and HLA genes. MAF of SNPs at the HLA and peri-HLA genes is higher compared to other genes.

**Figure 3.**
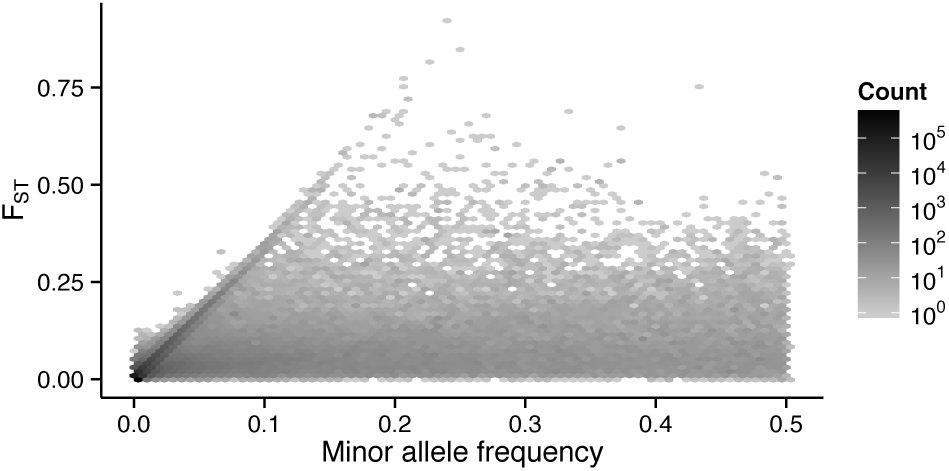
Population differentiation, measured by F_ST_, as a function of minor allele frequency at biallelic exonic SNPs from the 1000 Genomes Project phase 3 data.

Constraints imposed by allele frequencies on F_ST_ have been a topic of recent investigation (Roesti *et al.* 2012; Maruki *et al.* 2012; Elhaik 2012; Jakobsson *et al.* 2013; Edge and Rosenberg 2014; Alcala and Rosenberg 2017), and it has been shown that SNPs with very low minor allele frequencies (MAF) are bounded to low F_ST_ values. This relationship between MAF and F_ST_ is empirically illustrated for the 1000 Genomes exome data in Figure 3, which shows that F_ST_ is constrained to low values mainly in the range of low MAF (up to ~ 0.1 in the 1000G dataset). When MAF is above this value the constraint is no longer evident. In File S1, we analytically show the relationship between MAF and the maximum possible value of F_ST_.

This suggests that the large number of rare variants in the 1000G dataset, and the relative paucity of low MAF variants in the HLA SNPs could generate the observed differences in F_ST_ distributions between those groups (Figure 1).

To account for the effects of differences in SFS between HLA and control SNPs when contrasting population differentiation among those groups, we compared the F_ST_ of HLA and control SNPs within bins of MAF values (Figure 4). Contrary to what we observed without controlling for MAF, we now find that HLA and peri-HLA SNPs have significantly lower F_ST_ than at the control SNPs (Mann-Whitney two-tailed test p-value < 10^-5^ for all bins of MAF > 0.01). The bin with MAF < 0.01 shows a similar pattern when further split into smaller bins of MAF (Figure S3). Figure S4 shows FST distributions including outliers.

**Figure 4.**
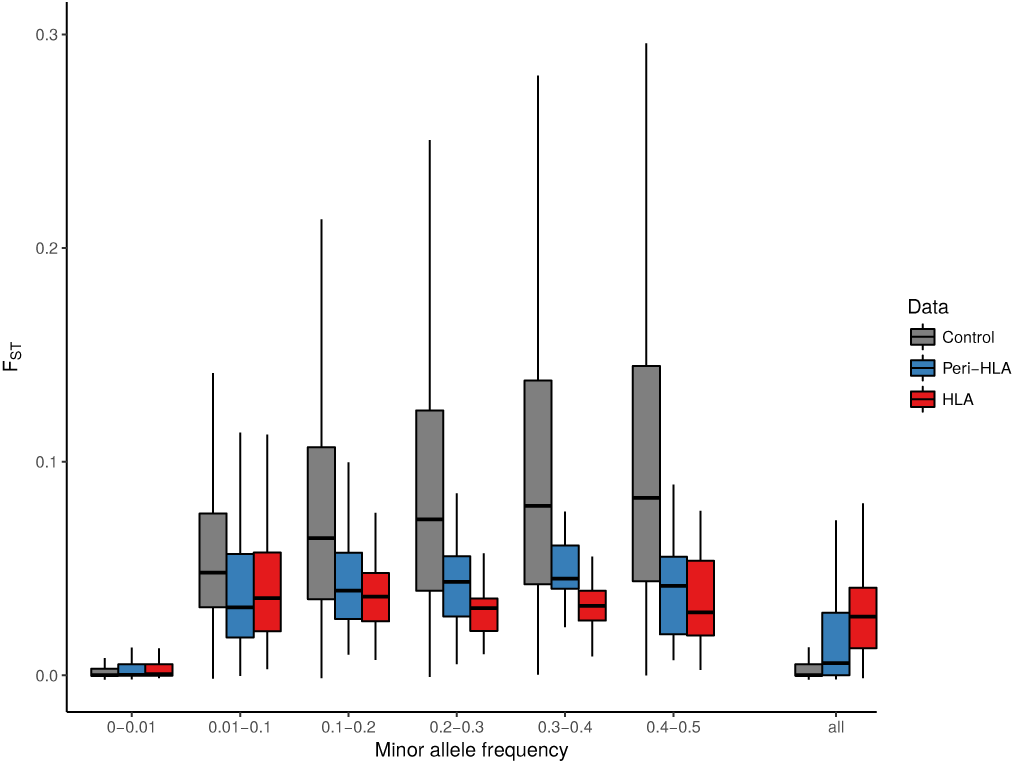
F_ST_ distributions per minor allele frequency (MAF) bin. HLA and Peri-HLA SNPs show lower F_ST_ than control SNPs in all bins with MAF > 0.01. Outliers were removed from figure, but not from statistical test, for better visualization. Figure S4 shows F_ST_ distributions including outliers.

This approach used SNPs outside the HLA and peri-HLA genes as controls for the HLA SNPs. However, because the HLA SNPs are located in 5 genes, they are not independent, due to both intra and inter-locus associations. As a consequence, our p-values could be inflated by treating a set of correlated SNPs as independent, and comparing them to a set of control SNPs which are in their majority independent. We controlled for this non-independence using a resampling approach, in which sets of linked SNPs were sampled from our control group, and their F_ST_ values were compared to those from the HLA SNPs, at each MAF bin (see Materials and Methods). After controlling for the non-independence of HLA SNPs, we confirmed that F_ST_ at HLA SNPs was significantly lower than at the resampled SNPs, at all bins of MAF higher than 0.01 (Table S4).

Another strategy to control for the constraint of MAF on F_ST_ is to estimate F_ST_ for multiple loci by computing a "ratio of the averages" of the numerator and denominator of the AMOVA-based F_ST_ estimator (see Methods). Reynolds *et al.* (1983) showed that this is the least biased estimate of average F_ST_ across multiple loci. The alternative, *i.e.,* computing the average F_STS_ for individual SNPs (an "average of ratios") creates a bias leading to an underestimation of F_ST_, the effect of the bias being more pronounced the more rare variants there is in the dataset (Bhatia *et al.* 2013). The ratio of averages approach, on the other hand, proportionally downweights the contribution of variants with low MAF. This results in a higher overall F_ST_ values for datasets rich in rare variants.

We explored how these different averaging methods impact the F_ST_ at HLA genes, and found that the "ratio of averages" approach (which controls for MAF) results in lower average F_ST_ at the HLA SNPs (0.04) than in the control SNPs (0.09). In stark contrast, using the average of individual loci F_ST_, we found higher F_ST_ values for the HLA SNPs (0.03) than genome-wide (0.01), as in our initial analysis that did not account for MAF. This further emphasizes the importance of accounting for the differences between the site frequency spectrum of HLA and control SNPs when assessing population differentiation.

### HLA-DP genes

The classical *HLA-DPA1* and *-DPB1* genes were excluded from the previous analysis because they show weak or no evidence of balancing selection (Solberg *et al.* 2008; Begovich *et al.* 2001), and some evidence of directional selection (supported by the observation that within individual populations a small number of alleles are present at a high frequency) (Hollenbach *et al.* 2001).

Consistently with being under a different selective regime, *HLA-DPA1* and *-DPB1* show a pattern of population differentiation which is different from the other classical HLA loci: F_ST_ at these genes is higher than in the control SNPs, even when minor allele frequency is controlled for (Figure S1).

### Contrasting F_ST_ of HLA SNPs and haplotypes

Next, we explored population differentiation at the haplotype level *(i.e.* with alleles defined by the coding sequence of each HLA gene). Haplotype level analyses were motivated by the idea that the fitness of individuals is more likely to be determined by the combination of SNPs they carry in a gene, rather than by individual SNPs, since it is the combination of SNPs that determines the peptides which HLA molecules present.

To compare F_ST_ for HLA haplotypes to a null distribution, we recoded each HLA gene as a biallelic locus (see Materials and Methods). Population differentiation at the recoded HLA haplotypes was then compared to differentiation at control SNPs and HLA SNPs, while controlling for minor allele frequency, as was done for SNPs.

The boxplots show that differentiation at HLA haplotypes and HLA SNPs are quite similar (Figure 5) and differentiation at HLA haplotypes is not significantly different from that of control SNPs, except for the MAF bin between 0 and 0.08. Thus, despite the existence of haplotypes which are specific to certain world regions, when the average MAF is considered and global F_ST_ is quantified, the degree of differentiation of HLA haplotypes is lower than that of control SNPs, as was found for HLA SNPs.

**Figure 5.**
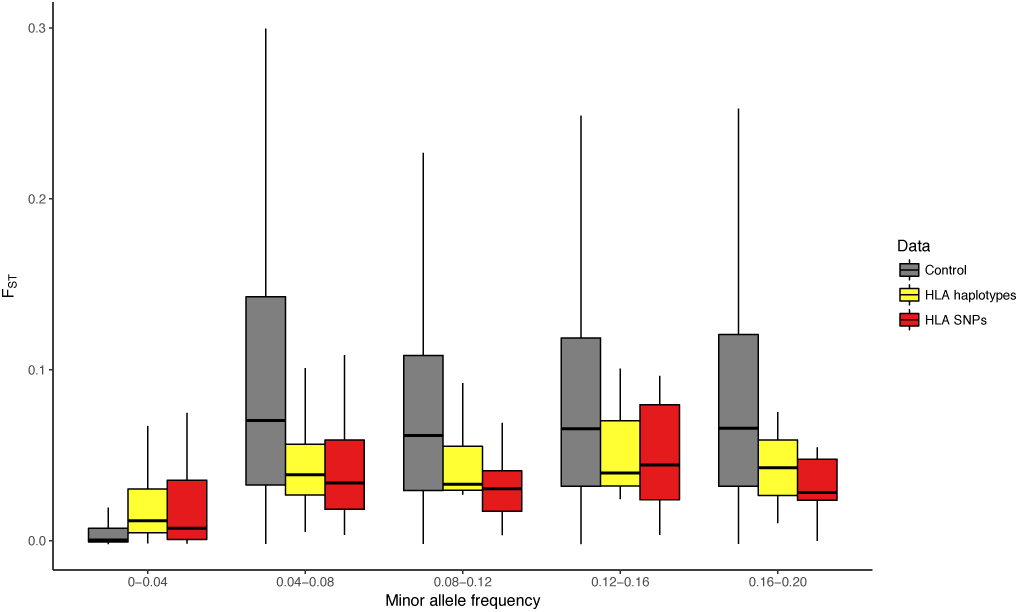
Population differentiation (F_ST_) as a function of minor allele frequency for the exonic control SNPs (gray), for exonic HLA SNPs (red) and for the haplotype-level *HLA* alleles (yellow) after recoding as biallelic (see Materials and Methods). No haplotypes had frequency higher than 0.2, therefore MAF bins were redefined.

It is worth noting that ambiguous HLA haplotype calls (i.e., instances where the typing method provided a set of possible allele calls) was resolved by choosing the assignment that minimized population differentiation. Thus, a more reliable assessment of haplotype-level differentiation will require less ambiguous haplotype calls.

### F_ST_ at HLA SNPs depends on divergence times

Our previous analyses examined global F_ST_, which captures patterns of differentiation among all 20 populations retained from the 1000 genomes full dataset. Next, we asked how specific populations contributed to our findings. In order to investigate this question, and to evaluate how the geographical scale (within and among continents) influences differentiation at the HLA, we analyzed F_ST_ between all pairs of populations.

We found that the lower differentiation at HLA SNPs as compared to control SNPs seen in our previous results (Figure 4) in seen for highly diverged populations (Figure 6) (i.e., contrasts involving populations from different continents).

**Figure 6.**
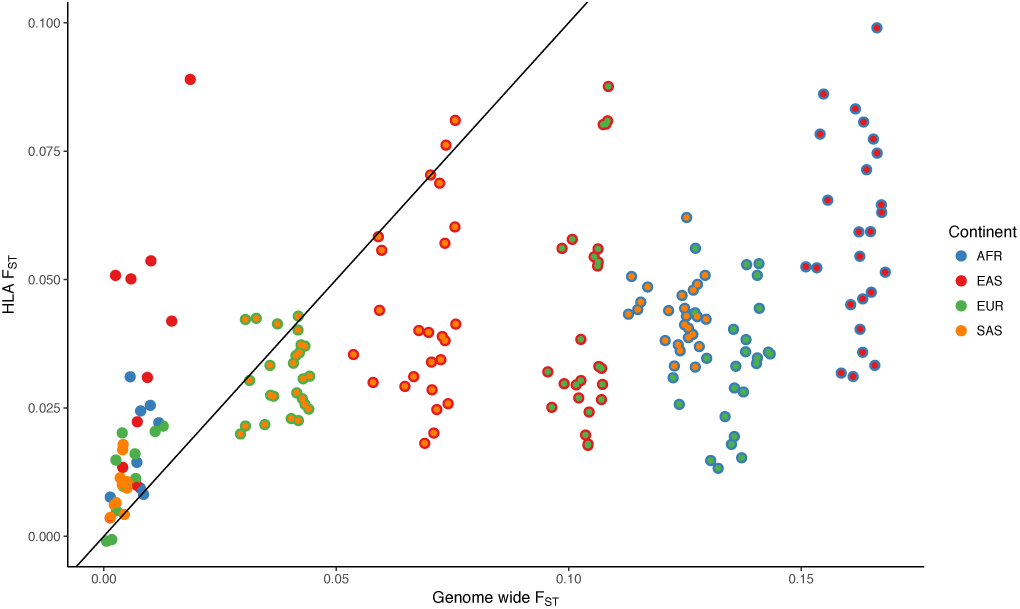
Average F_ST_ at the HLA and at the control SNPs for each pair of populations. Points with a single color represent pairs of populations from the same continent.

However, population pairs within the same continent show higher differentiation at the HLA SNPs compared to control SNPs. This effect was also observed in an independent analysis performed on the populations from the HGDP SNP dataset, with F_SC_ values showing a peak in the MHC region (indicating increased differentiation among populations within continents) (Figure S5).

## Discussion

### Population differentiation at HLA SNPs

In an overall analysis of F_ST_ among worldwide populations, we found significantly decreased genetic differentiation at HLA SNPs. We have shown that this result is critically dependent on the use of methods which appropriately account for the properties of the site frequency spectrum of HLA genes.

The decreased differentiation at HLA genes counters the expectation of a model of adaptation to local pathogens driving differentiation. However, we found that the overall pattern of lower differentiation reveals a greater complexity when we compare populations with different divergence times.

### Mind the MAF

Our results highlight the importance of accounting for minor allele frequency (MAF) when comparing F_ST_ values for different sets of SNPs, as previously shown by others (Roesti *et al.* 2012; Maruki *et al.* 2012; Elhaik 2012; Jakobsson *et al.* 2013; Edge and Rosenberg 2014). Not accounting for MAF leads to an underestimation of population differentiation for sets of SNPs rich in rare variants. When comparing sets of SNPs from genomic regions with different MAF distributions, this may result in a misleading interpretation of the selective regime acting on each region.

Alcala and Rosenberg (2017) recently arrived at the same results for the upper boundary of F_ST_ as a function of allele frequency in biallelic markers as we present on Supplemental File S1. Their work presents a more general investigation of the constraints of the frequency of the most frequent allele on F_ST_, using simulations under different migration models. Alcala and Rosenberg (2017) discuss the effect of this constraint in reducing the power of outlier tests which use high F_ST_ as a signature of local positive selection. Here we emphasize the effect of this constraint on interpreting F_ST_ values at regions under balancing selection, where the depletion of rare variants leads to higher overall population differentiation than in other genomic regions. However, when variants with similar MAF are compared, population differentiation in the region under balancing selection is actually lower, as we show for SNPs within HLA genes.

While the constraint of MAF on F_ST_ strongly affects differentiation at regions under positive and balancing selection, it is less problematic in regions under purifying selection. Under this selective regime, both an enrichment at low frequency variants and low population differentiation are expected. Since population differentiation in low frequency variants is constrained to low values, the relationship between MAF and F_ST_ leads both signatures in the same direction. This effect has been demonstrated in *Drosophila melanogaster* (Jackson *et al.* 2014).

### Contrast to previous studies

Two classical HLA genes, *HLA-DPA1* and *HLA-DPB1,* were exceptions to our finding of lower population differentiation at HLA SNPs. SNPs in those genes showed higher population differentiation, even when MAF was accounted for (Figure S1).

A genomic scan performed by Bhatia *et al.* (2011) also found a SNP near a HLA-DP gene with unusually high population differentiation (rs2179915, which is 30kb from *HLA-DPA2).* Similarly, Barreiro *et al.* (2008) found unusually high F_ST_ at the *HLA-DPB2* locus. These results indicate that the HLA-DP genes are not under balancing selection, but rather under directional selection that differs among populations. The studies which identified selection at these loci were designed with an emphasis on the detection of extremely high differentiation, and thus did not capture the pattern of low differentiation which is characteristic of most HLA loci.

Hofer *et al.* (2012), on the other hand, used an approach where regions of adjacent SNPs with extreme F_ST_ were scanned for, and detected the *HLA-C* locus among these, with evidence of unusually low F_ST_.The method used by Hofer *et al.* (2012) evaluates F_ST_ as a function of the heterozygosity between populations, which for biallelic markers is equivalent to the correction for MAF we applied here. These results show that when a test is designed to account for the possibility of unusually low F_ST_ values, and when the effect of minor allele frequency (or heterozygosity) on F_ST_ is accounted for, a signature of low differentiation which would otherwise not be detected can be found.

### Divergence time effect

By taking advantage of multiple populations made available by the 1000 Genomes project, we also examined if the excess of low differentiation at HLA SNPs holds at all timescales of differentiation. Interestingly, we find that for population pairs with low divergence (those from the same continent) F_ST_ at HLA SNPs is equal to or higher than in the control SNPs. For highly diverged population pairs (those from different continents), we consistently find lower differentiation among HLA SNPs. This result shows that a specific set of SNPs may differ in how they deviate from the genomic background depending on the timescale of population divergence.

To understand the process driving the increased differentiation for recently diverged populations (Figure 6) we used a simulation approach. First, we simulated a scenario where an ancestral population is under symmetric overdominant selection, and splits into two daughter populations, both under the same selective regime as the ancestor (details in Supplemental File S2). We refer to this scenario as "shared overdominance", and find that it results in differentiation between the daughter populations being reduced with respect to neutral expectations (Figure 7A and Figure 8A), in accordance with previous results (Schierup et al. 2000).

**Figure 7.**
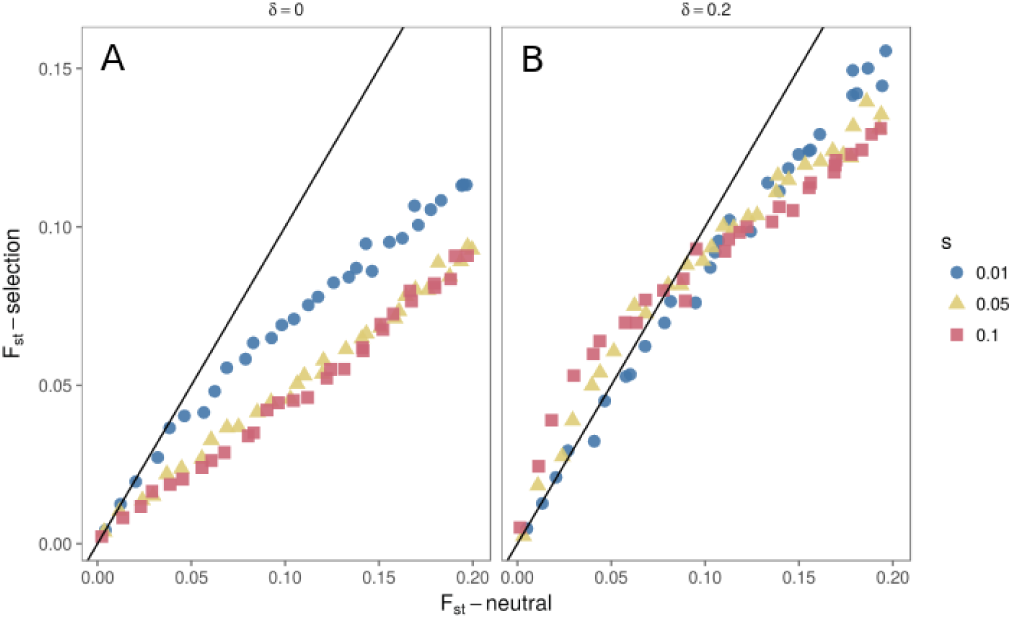
Relation between F_ST_ under neutrality and balancing selection. (A) daughter populations remain under the same regime of overdominance (shared overdominance) as the ancestral population. (B) One of the daughter populations experiences a shift in the fitness values (divergent overdominance), remaining under overdominance but with a new equilibrium value (changed by a value of *δ* = 0.2. In this case, for recent divergence times we find balancing selection can temporarily increase population differentiation, so long as selection is strong (s=0.05 or greater, *N*_*e*_ = 1000).

**Figure 8.**
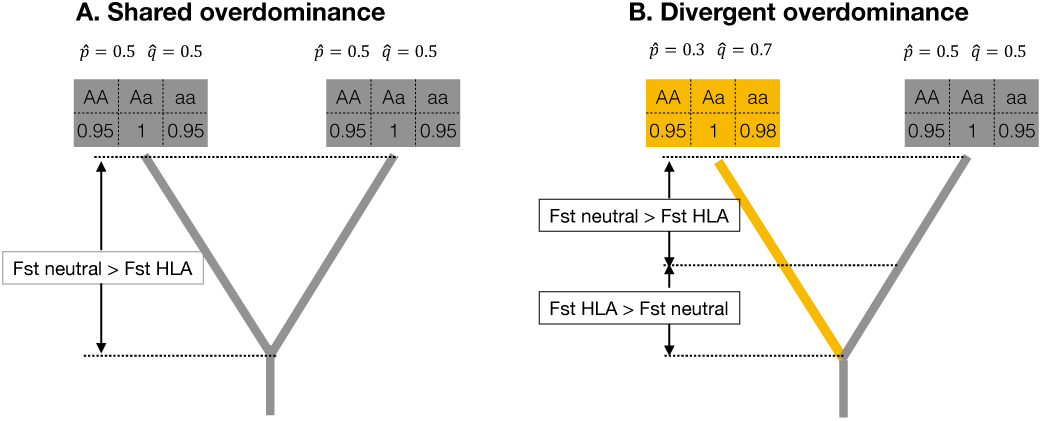
Schematic representation of HLA differentiation under two regimes of overdominance. Each tree represents a population pair experiencing selection according to fitness values presented in the boxes above the tips. In "shared over-dominance" both daughter populations share the same fitness values. In "divergent overdominance" the fitness values for one of the homozygotes changes in one of the daughter populations, even though the regime is still one of overdominance. In the divergent overdominance scenario, the equilibrium frequencies at the selected site differs among populations, and drives increased differentiation when divergence is recent (F_ST_ HLA > F_ST_ neutral). For the shared overdominance scenario F_ST_ neutral > F_ST_ HLA throughout the entire history of population divergence.

However, if the two populations remain under overdominant selection but differ for one of the homozygote fitness values (a scenario we refer to as "divergent overdominance"), differentiation can be increased with respect to neutrality for small divergence times (Figure 7B and Figure 8B). This result can be understood if we consider that the populations are extremely similar at the time of the split, so the effect of selection will be to favor changes in allele frequency between them (since homozygote fitness and therefore equilibrium frequencies of selected alleles will differ). As the two populations further diverge, the neutral sites will continue to diverge and will surpass the differentiation for the case of overdominance. Thus, by assuming that the fitness values of an overdominant model can change over time -which is highly plausible if we consider heterogeneity in pathogen populations affecting HLA fitness-increased F_ST_ at HLA for recently diverged populations can be explained.

Consistent with our results, recent studies have identified very recent adaptive change at HLA loci (Field *et al* 2016; Zhou *et al.* 2016), which could contribute to differentiation at a local scale without erasing signatures of long-term balancing selection.

## Acknowledgments

This research was financially supported by grants from São Paulo Research Foundation (FAPESP), The Brazilian National Council for Scientific and Technological Development (CNPq) and the Swiss National Science Foundation (SNSF). DYCB was funded by FAPESP scholarships #2012/22796-9 and #2013/12162-5, JC was funded by FAPESP scholarship #2015/19990-6 and DM has a FAPESP research grant #12/18010-0 and a CNPq productivity grant #308167/2012-0. JG was supported by grant 31003A-138180 of the SNSF.

